# Gut Microbiome Composition in Older Adults with PTSD Symptoms and Trauma-Exposed Controls: Findings from the GUMP Study

**DOI:** 10.64898/2026.08.23.746587

**Authors:** Cara M. Tout, Fiona Randall, Trish Wilson, Thomas Pace, Hattie Wright, Sophie C. Andrews, Mark Holmes, Bonnie L. Quigley

## Abstract

**Background:** Emerging evidence suggests that alterations in the gut microbiome may contribute to mental health outcomes through the gut-brain axis. However, older adults with post-traumatic stress disorder (PTSD) remain underrepresented in microbiome research. This pilot study investigated associations between PTSD symptoms, dietary fibre intake, cognitive function, and gut microbiome functional capacity in adults aged 50 years and older.

**Methods:** Participants with PTSD symptoms and trauma-exposed controls (TEC) completed validated assessments of mental health, trauma exposure, and dietary fibre intake. A subset of participants provided stool samples for microbiome analysis and undertook cognitive function assessment. Quantitative PCR was used to assess phylum-level taxonomy and butyrate-producing bacterial pathways (terminal butyrate generating enzyme), with abundances normalised to the 16S rRNA gene.

**Results:** Participants with PTSD demonstrated significantly greater mental health symptom burden and poorer performance on cognitive tasks related to executive function, working memory, and learning. Dietary fibre intake did not differ significantly between PTSD and TEC groups and no significant differences in overall microbial composition were identified at the phylum level. In contrast, differences were more apparent when assessing functional microbiome pathways, with butyrate kinase abundance significantly lower in PTSD participants than TEC participants. When stratified by fibre intake, a greater butyrogenic capacity was observed in the High fibre TEC participants compared to the Low fibre TEC participants, while little difference was observed in the PTSD fibre-stratified groups. Substantial inter-individual variation was also evident across both taxonomic and functional measures.

**Conclusions:** These findings suggest that functional characteristics of the gut microbiome may provide greater insight into PTSD-related biological processes than broad taxonomic measures alone. Dietary fibre intake may be associated with greater butyrate-producing capacity in trauma-exposed older adults without PTSD symptoms, although this relationship appeared less evident among older adults living with PTSD symptoms. These findings support further investigation of microbiome function, diet, and cognition within the gut-brain axis. Larger studies incorporating metagenomic and metabolomic approaches are warranted.

## Introduction

### Post-Traumatic Stress Disorder (PTSD)

Up to 70% of adults worldwide will experience trauma at some point in their lives (Kessler et al., 2017). Of Australian adults, 42% have reported experiencing an Adverse Childhood Experience (ACE) before turning eighteen, with childhood trauma exposure associated with an increased risk of chronic mental and physical health complications across the lifespan (Barrett et al., 2026). Though the majority of the population experiences trauma, only approximately 5.6-10% will go on to develop Post-Traumatic Stress Disorder (PTSD), depending on the population and type of trauma exposure (Kessler et al., 2017; Zohar et al., 2000).

PTSD is currently classified in the Diagnostic and Statistical Manual of Mental Disorders V (DSM-5) as a Trauma- and Stressor-Related Disorder (American Psychiatric Association, 2013). It is comprised of four symptom clusters: intrusion, avoidance, increased arousal and reactivity, and negative mood and cognition (American Psychiatric Association, 2013). Those with PTSD frequently exhibit comorbid mental health conditions, such as major depressive disorder, anxiety disorders, and substance use disorders (Barrett et al., 2026; Kessler et al., 2017). Furthermore, approximately half of those with PTSD will experience chronic symptom presentation (Steinert et al., 2015), highlighting the chronic and debilitating nature of this disorder and the significant global health burden it presents.

### PTSD, Cognition, and Ageing

Despite a lower 12-month PTSD prevalence in older adults (2.6-3.0%) compared to younger and middle-aged adults (4.3-5.2%) (Kessler et al., 2005; Reynolds et al., 2016), older adults with PTSD remain at increased risk of disability, accelerated ageing and related morbidity, and cognitive decline, including an elevated risk of dementia (Byers et al., 2014; Yaffe et al., 2010). Consequently, older adults with PTSD represent an underrepresented population for investigating the long-term neurophysiological effects of trauma. PTSD is associated with memory impairment and executive dysfunction, which overlap with cognitive changes seen in dementia and are linked to increased risk of cognitive decline (Sumner et al., 2017). Older adults with chronic, complex, or untreated PTSD therefore represent a key at-risk cohort. Neurological dysregulation associated with PTSD has physiological impacts across multiple body systems, including the circulatory and gastrointestinal systems. Notably, those with PTSD are at increased risk for developing cardiovascular disease (CVD) (Beristianos et al., 2016), gastrointestinal disorders such as Inflammatory Bowel Syndrome (IBS) (Taft et al., 2019), and dementia (Nilaweera et al., 2020).

### The Gut Microbiome

Chronic stress associated with PTSD has been shown to disrupt the gut microbiome (Hemmings et al., 2017; Leclercq et al., 2016; Malan-Muller et al., 2022), a complex ecosystem of commensal bacteria involved in neurotransmitter production, immune regulation, and gastrointestinal health. This suggests that the gut microbiome may reflect systemic dysfunction in PTSD, offering potential for novel biomarker development (Zeamer et al., 2023).

Microbial profiling has revealed alterations in the abundance of prominent bacterial taxa, as well as specific bacterial genera, in individuals with PTSD (Malan-Muller et al., 2022). For example, Bajaj et al. (2019) found that individuals with PTSD had decreased abundance of *Ruminococcaceae* and *Lachnospiraceae* (both from the major phylum *Firmicutes*). Multiple studies have also identified changes in the lower-abundance phyla, including increased abundance *of gamma-Proteobacteria* in combination with decreased *Actinobacteria* and *Verrucomicrobia* abundance during PTSD (Geier et al., 2024; Marano et al., 2026). PTSD symptom severity, increased hyperarousal, and symptom load have all been associated with microbiome dysbiosis characterised by reduced abundance of these smaller phyla, as well (Hemmings et al., 2017).

### Dietary Fibre and Butyrate

Dietary fibre is a food that is resistant to digestion by human enzymes and reaches the colon largely intact, where it acts as a substrate for microbial fermentation (Maier et al., 2017). As a major energy source for commensal gut bacteria, dietary fibre influences microbiome composition, metabolism, and short-chain fatty acid production, with microbial taxa exhibiting preferences for specific fibre sources (Gill et al., 2021; Sankarganesh et al., 2025). Fermentation of dietary fibre produces short-chain fatty acids (SCFAs), which contribute to intestinal barrier integrity, immune regulation, and metabolic health. Dietary fibre therefore represents a key environmental factor influencing gut microbiome composition and metabolite production. SCFAs are microbial metabolites produced by commensal gut bacteria (Fusco et al., 2023). The three major SCFAs are acetate, propionate, and butyrate (produced in a ratio of 60:20:20, respectively) (Topping & Clifton, 2001). Butyrogenic bacteria rely on enzymes from the CoA-transferase (CoAT) gene superfamily, containing either the butyryl-CoA:acetate CoA-transferase (*but*) gene or the butyrate kinase (*buk*) gene as the terminal enzyme in the pathway to generate butyrate (Singh et al., 2023; Vital et al., 2015). Variation in the abundance of these pathways may reflect functional differences within the gut microbiome and dietary patterns (Van-Wehle & Vital, 2024; Vital et al., 2014).

### Study Hypotheses

Given the known links between PTSD, comorbid mental health conditions and cognitive performance, this study hypothesised that participants living with PTSD symptoms would demonstrate a higher mental health symptom burden and poorer cognitive performance than TEC participants. Additionally, based on established links between dietary fibre, the gut microbiome and PTSD, this study further hypothesised that PTSD symptoms would be associated with changes in gut microbiome composition and butyrate-production pathways, with higher dietary fibre intake being associated with greater butyrate-producing capacity.

## Methods

### Study Design and Participants

The <u>Gu</u>t <u>M</u>icrobiome in <u>P</u>TSD in older adults (GUMP) study was a two-phase cross-sectional study. Participants aged ≥50 years were recruited from the Sunshine Coast region (Queensland, Australia) through university, community, and word-of-mouth advertising.

Participants were excluded if they reported following a specialised diet (e.g., vegetarian, vegan, or gluten-free) or had used antibiotics within the previous month due to their known effects on gut microbiome composition (Elvers et al., 2020; Fackelmann et al., 2025; Tomova et al., 2019). Participants selected for biological sampling were chosen to make PTSD symptom or control groups and these groups were matched for age, sex, and estimated dietary fibre intake. Ethical approval was granted by the UniSC Human Research Ethics Committee (A252734).

### Part 1: Online Surveys

Participants provided electronic informed consent and completed a set of validated questionnaires via Qualtrics. Questionnaires assessed dietary fibre intake (FibreScreen; (Rijnaarts et al., 2021)), symptoms of anxiety, depression, and stress (Hospital Anxiety and Depression Scale [HADS] (Zigmond & Snaith, 1983); Depression Anxiety Stress Scale [DASS-21] (Lovibond & Lovibond, 2011)), trauma exposure (Life Events Checklist for DSM- 5 [LEC-5] (Weathers et al., 2013)), and PTSD symptom severity (PTSD Checklist for DSM-5 [PCL-5] (Blevins et al., 2015)). Participants meeting DSM-5 symptom cluster criteria and scoring ≥33 on the PCL-5 were classified as probable PTSD and were assigned into the PTSD symptom group, while trauma-exposed participants not meeting PTSD criteria were classified as trauma-exposed controls (TEC). Participants were classified into low- (<25 g/day) and high-fibre (≥25 g/day) groups according to the Australian Adequate Intake recommendation for dietary fibre based on FibreScreen results.

### Part 2: Biological Sampling and Cognitive Assessment

As subset of eligible participants attended a study visit at the Thompson Institute, during which an at-home faecal collection kit was returned, and fasting capillary blood samples and cognitive assessments were collected. All data and biological samples were assigned de-identified study codes.

Participants collected faecal samples at home using BunnyWipe collection kits (Zymo Research, Australia) containing DNA/RNA Shield preservative allowing room temperature storage until return to the study team. Returned samples were then stored at 4°C until DNA extraction.

Fasting capillary blood samples were collected by finger prick using sterile lancets and mini- serum separator tubes. Blood samples were allowed to clot for 30 minutes at room temperature before centrifugation at 2,000 × g for 5 minutes. Serum was aliquoted and stored at −80°C until analysis.

Cognitive function was assessed using the Cambridge Neuropsychological Test Automated Battery (CANTAB) (Robbins et al., 1994). Six tasks were administered that assessed motor function (MOT), visual learning and memory (PAL, DMS), psychomotor speed (RTI), spatial working memory (SWM), and sustained attention (RVP). Outcome measures demonstrating appropriate variation across participants were selected for subsequent analyses (Abbott et al., 2018).

### DNA Extraction

Microbial DNA was extracted from faecal samples using the Zymo Quick DNA Faecal Microbe Miniprep Kit (Zymo Research, Australia) according to the manufacturer’s instructions. DNA concentration was assessed using a NanoDrop Microvolume Spectrophotometer (Thermo Fisher Scientific, Australia), and purified DNA was stored at 4°C until analysis to avoid repeated freeze-thaw cycles.

### Microbiome Analysis

Gut microbiome profiling was conducted using qPCR to assess both taxonomic composition and functional butyrate-production pathways. Taxonomic profiling targeted total 16S gene copy number (Universal), *Firmicutes, Bacteroidetes, Actinobacteria, gamma-Proteobacteria,* and *Verrucomicrobia,* while functional profiling targeted the butyryl-CoA:acetate CoA-transferase (*but*) and butyrate kinase (*buk*) genes. While the *buk* gene from the gut microbiome could be quantified using a single qPCR assay (Vital et al., 2013), the *but* gene required four separate assays to capture and quantify the genetic diversity present in this community (subgroups A, C, D and F) (Daskova et al., 2021).

Primer sets were selected from published studies (De Gregoris et al., 2011; Daskova et al., 2021; Vital et al., 2013; Yang et al., 2015) and optimised for use with participant faecal DNA. Primer sequences are presented in Supplementary Table 1. All qPCR assays were performed in 20 μL reaction volumes using Platinum SuperFi II PCR Mastermix (Invitrogen, Thermo Fisher Scientific, Australia), target-specific primers, template DNA, and deionised water. Primer-specific concentrations for qPCR assays are listed in Supplementary Table 2. No-template controls (NTCs) were included in each assay. Standard curves were generated for each target and used to assess assay performance and calculate starting quantity (SQ) values. Assay conditions are listed in Supplementary Table 3. Only assays demonstrating acceptable performance (R² > 0.950) were included in subsequent analyses.

### Relative Abundance Calculations

Relative abundance was calculated by normalising target SQ values to the universal 16S rRNA gene target (target SQ/universal SQ). Relative abundance values were interpreted as estimates of microbial presence rather than exact measures of bacterial abundance due to variation in 16S rRNA gene copy number among taxa.

### Statistical Analysis

All statistical analyses were performed in RStudio (2025). Data visualisation was conducted using ggplot2 (Wickham, 2016), while data manipulation and statistical analyses were performed using tidyverse packages (Wickham et al., 2019). Continuous variables were assessed for normality using the Shapiro-Wilk test. Independent-samples t-tests or Welch’s t-tests were used for normally distributed variables, while non-parametric data were analysed using Wilcoxon rank-sum or Kruskal-Wallis tests, as appropriate. Associations between microbial abundance, dietary variables, and clinical measures were assessed using Spearman’s rank correlation coefficient (ρ). Categorical variables were compared using Chi-square tests of independence. Linear regression analyses included Cook’s distance analysis for influential observations and sensitivity analyses (excluding the most influential observation) resulting in findings that were robust to individual data points. Principal Component Analysis (PCA) was used as an exploratory visualisation technique to assess clustering and patterns of variation within microbiome and clinical datasets. Statistical significance was defined as p < 0.05.

## Results

### Participant Characteristics

Table 1 summarises the demographic and key clinical characteristics of participants in Part 1 and Part 2 of the study, between the two study cohorts (PTSD and TEC). As a note, participants self-selected for Part 1 (survey was open to the general public) but were selected and contacted by the research team for participation in Part 2 to match age, sex and dietary fibre intake between the PTSD and TEC study groups.

**Table 1.** Participant demographics and mental health characteristics.

| Characteristics | Part 1 Survey |  |  | Part 2 Biological Sampling |  |  |
| --- | --- | --- | --- | --- | --- | --- |
|  | TEC<br>(n = 81) | PTSD<br>(n = 29) | p-value | TEC<br>(n = 16) | PTSD<br>(n = 15) | p-value |
| Demographics |  |  |  |  |  |  |
| Sex | 14 M / 66 F | 5 M / 25 F | N/A | 3 M / 13 F | 2 M / 13 F | N/A |
| Age (years) | 63.4 ± 8.73 | 56.6 ± 6.61 | < 0.001 | 59.3 ± 6.17 | 58.6 ± 6.28 | 0.753 |
| Mental Health Measures |  |  |  |  |  |  |
| PCL-5 Score | 9.02 ± 8.98 | 52.2 ± 14.40 | < 0.001 | 9.75 ± 8.14 | 44.2 ± 12.8 | < 0.001 |
| HADS Anxiety Score | 5.03 ± 3.31 | 13.8 ± 4.81 | < 0.001 | 5.81 ± 3.15 | 12.4 ± 4.56 | < 0.001 |
| HADS Depression Score | 3.45 ± 2.98 | 11.1 ± 4.75 | < 0.001 | 3.25 ± 2.86 | 8.67 ± 3.87 | < 0.001 |
| DASS-21 Anxiety Score | 1.86 ± 2.20 | 10.40 ± 5.30 | < 0.001 | 2.0 ± 1.75 | 7.67 ± 5.29 | < 0.001 |
| DASS-21 Depression Score | 3.19 ± 3.21 | 13.12 ± 5.43 | < 0.001 | 3.12 ± 2.70 | 10.4 ± 5.10 | < 0.001 |
| DASS-21 Stress Score | 4.84 ± 3.39 | 13.08 ± 4.55 | < 0.001 | 5.88 ± 3.38 | 10.3 ± 3.64 | < 0.001 |
| Diet |  |  |  |  |  |  |
| Fibre Intake (g) | 23.6 ± 8.8 | 19.5 ± 10.1 | 0.056 | 24.73 ± 7.3 | 19.49 ± 10.3 | 0.116 |

\*Values are presented as mean ± standard deviation (range) or count, as appropriate. All participants reported at least one traumatic exposure. Normality was assessed using the Shapiro-Wilk test. Group comparisons were conducted using Welch's t-tests for normally distributed variables and Wilcoxon rank-sum tests for non-normally distributed variables.

All participants in Part 1 reported exposure to at least one traumatic event, and many reported multiple trauma types. The most commonly reported trauma exposures were accidents (70%, n = 77), illness or injury (69%, n = 76), natural disasters (68%, n = 75), physical assault (67%, n = 74), death exposure (57%, n = 63), sexual assault (42%, n = 47), and war/combat exposure (29%, n = 32).

In Part 1, mental health measures were strongly and positively correlated with one another (r_s_ = 0.73-0.86, all *p* < 0.001). The strongest association was observed between PCL-5 scores and DASS Stress scores (r_s_ = 0.86, *p* < 0.001), indicating that greater PTSD symptom severity was associated with higher perceived stress.

### Cognitive Function in Older Adults

Cognitive outcomes are presented in Table 2. PTSD participants demonstrated significantly poorer performance on measures of visual learning and memory (PALTEA), sustained attention latency (RVPMDL), and spatial working memory strategy (SWMS) compared with TEC participants (all *p* < 0.05). No significant differences were observed for sustained attention errors (RVPTM), delayed matching to sample performance (DMSPC), or spatial working memory errors (SWMBE) (Table 2).

**Table 2.** Cognitive test outcomes.

| CANTAB Test Variables | TEC ($n = 16$) | PTSD ($n = 15$) | $p$-value |
| --- | --- | --- | --- |
| Visual memory & new learning<br>(PALTEA, # errors, lower score better) | 15.9 ± 10.9 | 26.9 ± 13.9 | 0.019* |
| Sustained attention time<br>(RVPMDL, time in sec, faster time better) | 463.0 ± 49.8 s | 547.0 ± 99.9 s | 0.007* |
| Sustained attention errors<br>(RVPTM, total misses, lower score better) | 14.8 ± 8.6 | 21.1 ± 9.2 | 0.060 |
| Short-term & working memory<br>(DMSPC, % correct, higher % better) | 87.2 ± 10.5% | 87.3 ± 7.5% | 0.779 |
| Spatial working memory errors<br>(SWMBE, # errors, lower score better) | 12.6 ± 11.4 | 16.9 ± 6.4 | 0.276 |
| Spatial working memory strategy<br>(SWMS, search strategy score, lower is better) | 7.1 ± 2.5 | 9.3 ± 2.1 | 0.007* |

PTSD = living with PTSD symptoms, TEC = trauma-exposed control. Mean ± SD, test statistic is Wilcoxon rank sum for PALTEA, RVPMDL, DMSPC, SWMBE, SWMS, and a Welch's t-test for RVPTM (based on Shapiro-Wilk's test of normality).

As cognitive performance can be influenced by age independently of PTSD symptom severity, Spearman correlations between age, PCL-5 scores, and CANTAB outcomes were assessed across the Part 2 study group. Higher PCL-5 scores were significantly associated with less efficient spatial working memory strategy (SWMS; r_s_ = 0.54, *p* = 0.002, Figure 1A) and poorer sustained attention time (RVPMDL; r_s_ = 0.40, *p* = 0.024, Figure 1C). Interestingly, while age was not associated with spatial working memory strategy (SWMS; r_s_ = -0.1, *p* = 0.596, Figure 1B), it was significantly associated with poorer sustained attention time (RVPMDL; r_s_ = 0.55, *p* = 0.001, Figure 1D). No significant associations were observed between age or PCL-5 scores and sustained attention errors (RVPTM), visual memory and new learning (PALTEA), spatial working memory errors (SWMBE), or short-term and working memory (DMSPC) (all *p* > 0.05).

**Figure 1.**
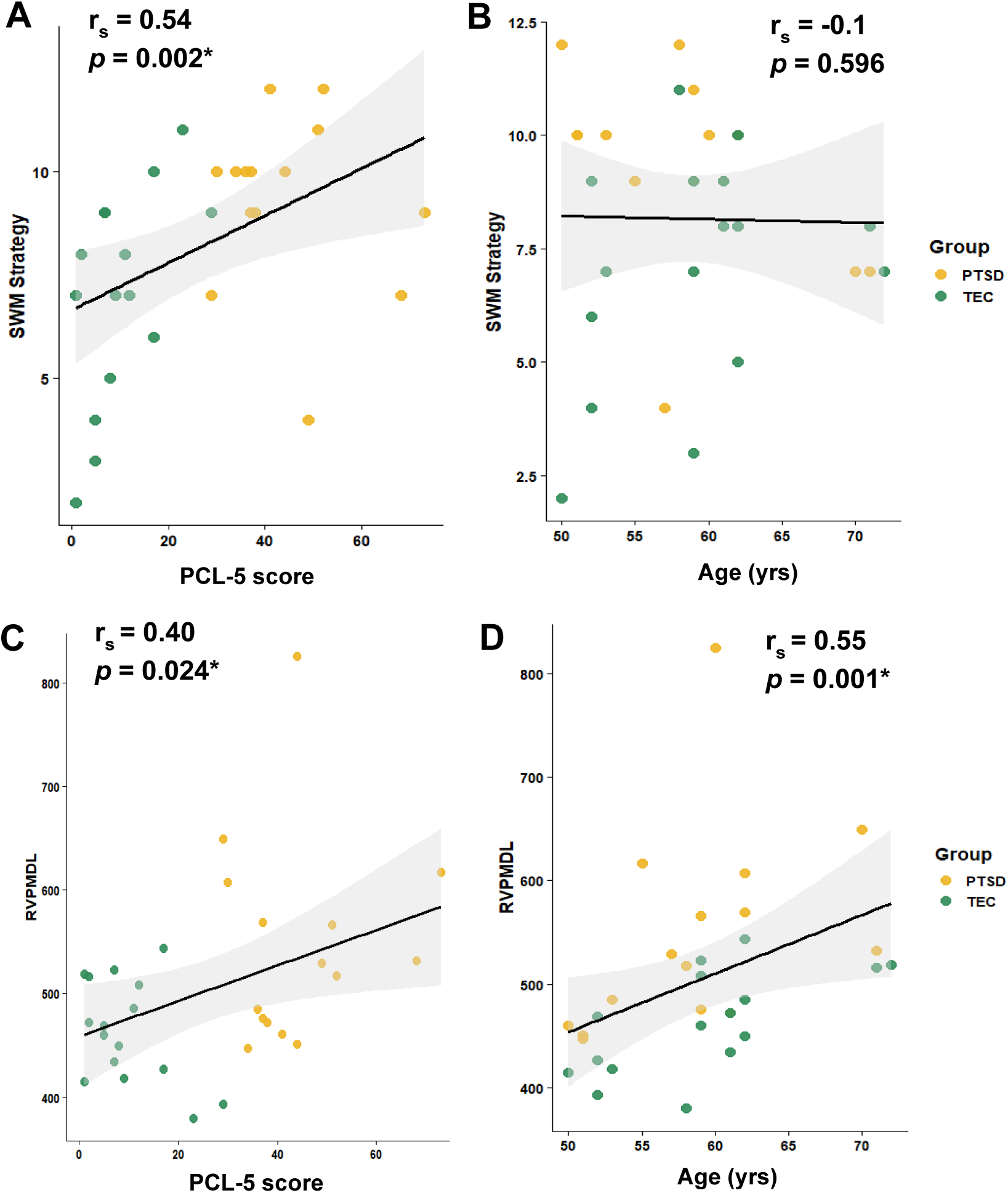
Associations between PTSD symptom severity, age, and selected cognitive performance measures. **(A)** Spatial working memory strategy (SWMS) and PCL-5 scores. Lower SWMS scores indicate better performance. **(B)** SWMS and age. **(C)** Sustained attention time (RVPMDL) and PCL-5 scores. Lower RVPMDL scores indicate better performance. **(D)** Age and RVPMDL.

Given the significant associations observed between PCL-5 scores and both SWMS and RVPMDL scores, linear regression analyses were performed to determine whether the associations between PTSD symptom severity and cognitive performance remained after adjustment for age. PTSD symptom severity remained significantly associated with SWMS (β = 0.057, *p* = 0.011) and RVPMDL (β = 1.883, *p* = 0.008) after accounting for age, while age itself was independently associated with RVPMDL (β = 6.210, *p* = 0.007) but not SWMS (β = 0.010, *p* = 0.887).

### Dietary Fibre Intake

During recruitment for Part 2 of this study, participants from each study group (TEC and PTSD) were matched as closely as possible on dietary fibre intake. Therefore, there was no significant difference in fibre intake between the two groups. Consistent with this matching approach, most dietary fibre categories did not differ significantly between the Part 2 TEC and PTSD groups, except for fibre obtained from fruit consumption, which did differ significantly between groups (*p* < 0.05).

### Phylum-Level Composition of the Gut Microbiome

For microbiome analysis, PTSD and TEC study groups were further subdivided into Low or High groups based on fibre intake (less or more than the recommended 25 g of fibre/day based Australian dietary guidelines, respectively). No significant differences in phylum relative abundance (*Firmicutes, Bacteroidota, Actinobacteriota, gamma-Proteobacteria*, or *Verrucomicrobiota*) were observed between the TEC Low, TEC High, PTSD Low, and PTSD High groups (all *p* > 0.05; Supplementary Table 4). *Firmicutes* and *Bacteroidota* were the dominant phyla across all participants, consistent with published human gut microbiome profiles (Cunningham et al., 2021).

### Butyrate Synthesis Capacity of the Gut Microbiome

Relative abundances of butyrate-producing pathways are presented in detail in Supplementary Table 5. No significant differences were observed between TEC and PTSD participants for the *but* gene or any of its four subgroups (all *p* > 0.05). In contrast, *buk* gene abundance was significantly lower in PTSD participants than TEC participants (Figure 2A). While considerable inter-individual variation was observed in *buk* gene abundance, higher *buk* gene abundances appeared to cluster within the TEC high-fibre group, although this difference across fibre-stratified study groups was not statistically significant (Kruskal-Wallis p > 0.05) (Figure 2A). To further explore the relationship between PTSD symptom severity and *buk* abundance, linear regression analyses were performed using log_10_-transformed *buk* abundance as the outcome variable. Higher PCL-5 scores were significantly associated with lower *buk* gene abundance (β = -0.017, p = 0.027) (Figure 2B). Interestingly, fibre intake was not significantly associated with *buk* gene abundance (β = 0.019, *p* = 0.251).

**Figure 2.**
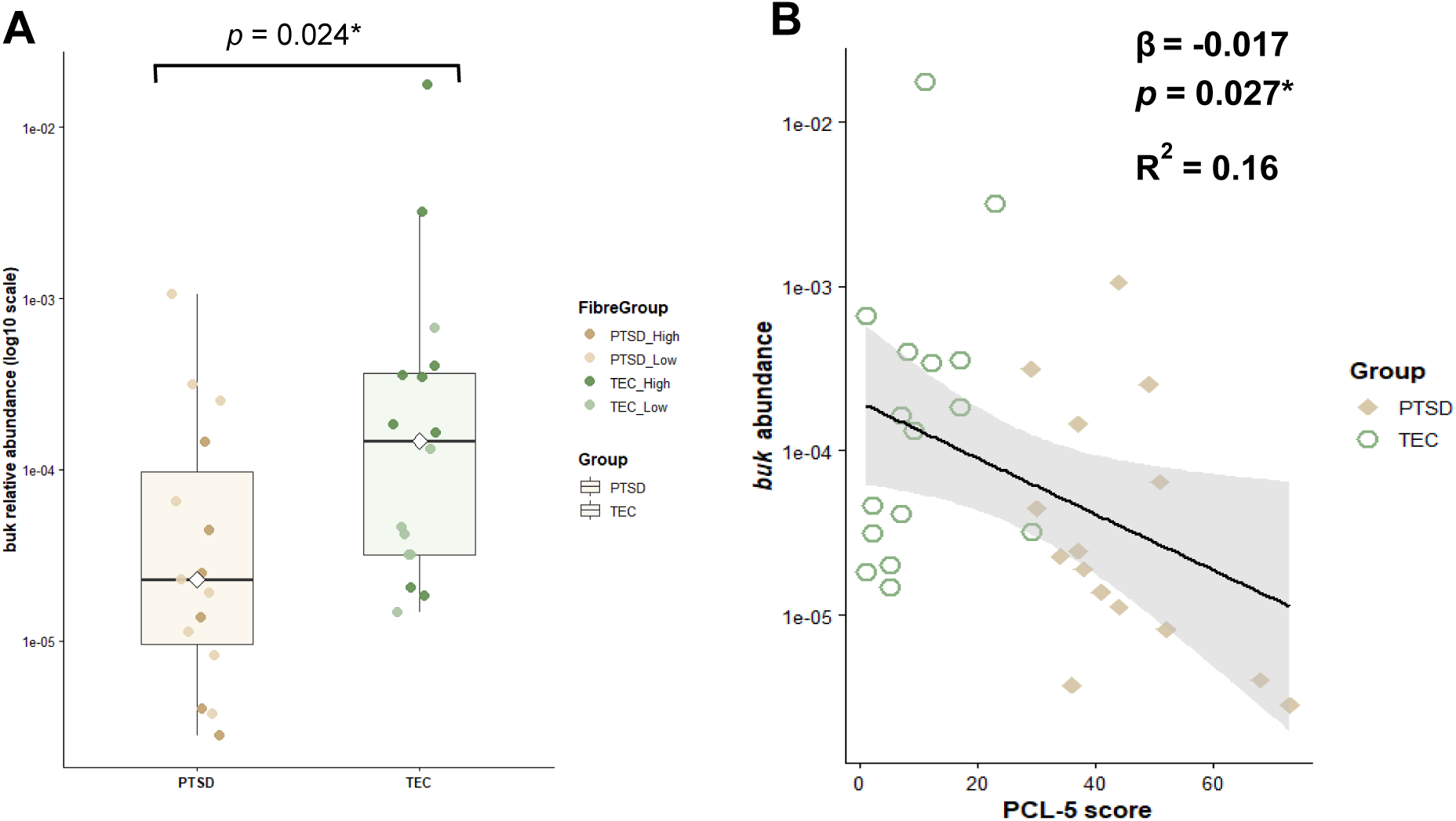
Relative *buk* abundance and its association with PTSD symptom severity. **(A)** Relative *buk* gene abundance by TEC and PTSD participants and fibre intake. **(B)** Relationship between PTSD symptom severity (PCL-5 score) and relative *buk* abundance. The solid line represents the fitted linear regression with 95% confidence interval (grey envelope).

When total butyragenic capacity from combined *but* and *buk* gene abundances was considered, the TEC study group showed an overall trend towards more butyragenic capacity compared to the PTSD symptom group, though this difference was not statistically significant (Figure 3A, Supplementary Figure 1). To further explore the magnitude of fibre-associated differences within each study group, Hedges’ *g* effect sizes were calculated for the High versus Low fibre comparison separately within the PTSD and TEC cohorts (Figure 3B). A negligible effect was observed within the PTSD cohort (Hedges’ *g* = -0.18), while a moderate effect was observed within the TEC cohort (Hedges’ *g* = 0.59). Although confidence intervals were wide, these findings suggest a greater difference in total butyrate abundance between High and Low fibre participants in the TEC cohort than in the PTSD cohort.

**Figure 3.**
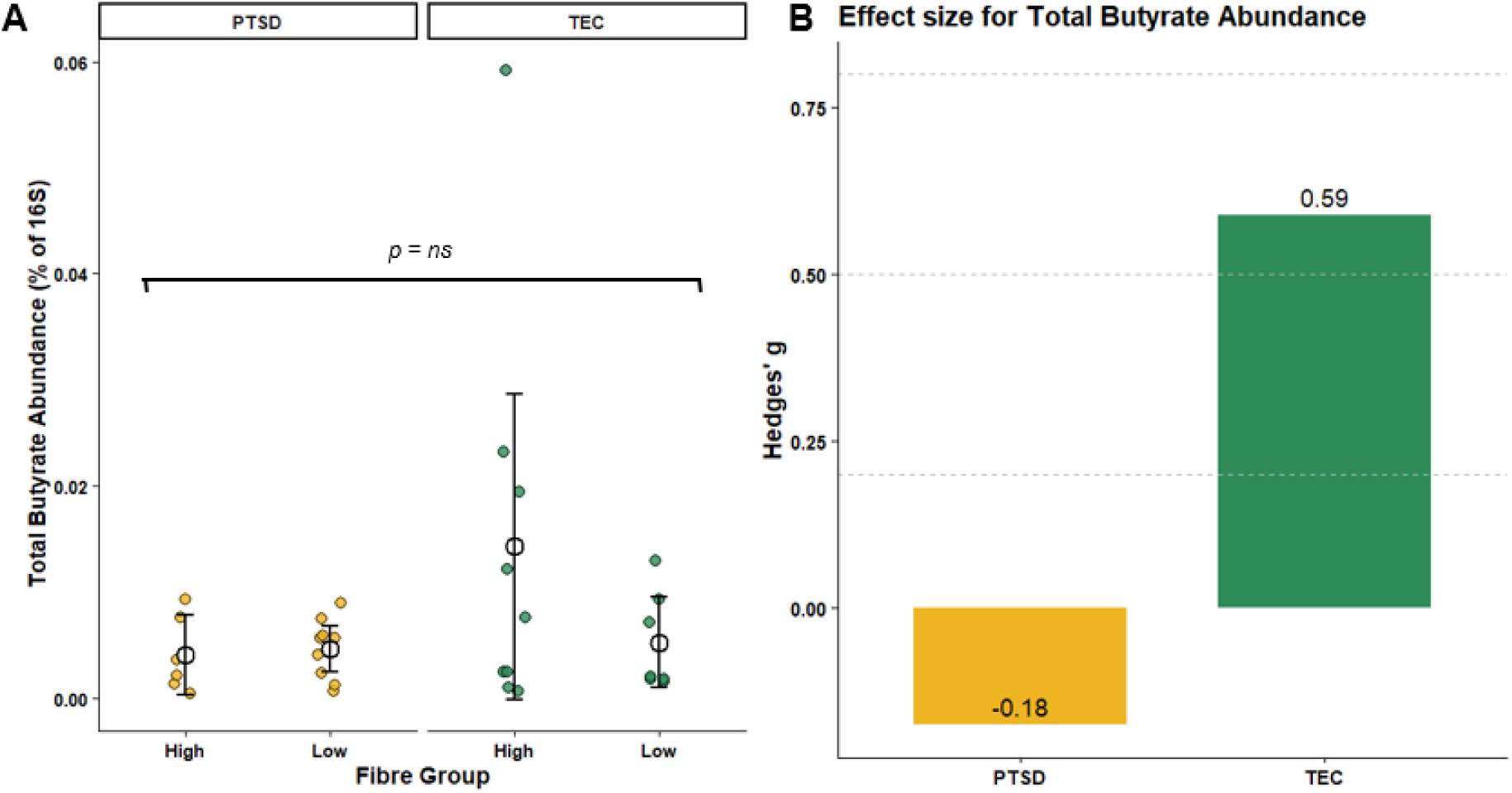
Comparison of total butyrate abundance (combined *but* cluster and *buk* gene abundance) across study groups and fibre intake categories. **(A)** Individual participant total butyrate abundance stratified by study group and fibre intake. **(B)** Hedges’ g effect sizes representing the magnitude of the High versus Low fibre group difference in total butyrate abundance, calculated separately for the PTSD and TEC cohorts.

Inspection of model diagnostics identified one influential observation within the TEC cohort (Cook’s distance = 0.68) (Figure 3A). Sensitivity analyses excluding this observation reduced the estimated effect size for the fibre group comparison from Hedges’ *g* = 0.59 (95% CI: -0.38 to 1.54) to g = 0.45 (95% CI: -0.53 to 1.41); however, the direction and overall interpretation of the result remained unchanged. In both analyses, total butyrate abundance was higher in the high-fibre TEC group compared with the low-fibre group, suggesting that the observed pattern was not driven solely by this participant (Figure 3B). Additionally, substantial inter-individual variability in butyrate pathway abundance was observed across participants (Supplementary Figure 1). While such variability is expected in microbiome analyses, the combination of high between-participant heterogeneity and small subgroup sample sizes likely contributed to the wide confidence intervals and limited ability to detect statistically significant differences.

## Discussion

This study had two main hypotheses: 1) participants living with PTSD symptoms would demonstrate a higher mental health symptom burden and poorer cognitive performance than TEC participants and 2) that PTSD symptoms would be associated with changes in gut microbiome composition and butyrate-production pathways, with higher dietary fibre intake being associated with greater butyrate-producing capacity.

Our results supported the first hypothesis, with participants in the PTSD study group presenting with significantly greater mental health symptom burden across multiple validated mental health scales (HADS, DASS-21, PCL-5), as well as demonstrating poorer performance on tasks assessing executive function, spatial working memory, and learning. This was despite the fact that dietary fibre intake did not differ significantly between the PTSD and TEC study groups.

Regarding our second hypothesis, no significant differences in overall microbiome composition were observed between PTSD and TEC participants at the phylum level, although differences emerged when assessing functional butyrate-production pathways. By specific butyrate generating pathway, *buk* gene abundance was significantly lower in PTSD participants than TEC participants. When total butyrate-producing capacity was estimated by using the combined abundance of all *but*-clusters and *buk*, greater butyrogenic capacity was observed in the High fibre TEC participants when compared to their Low fibre counterparts. However, little difference was observed between the PTSD High and Low fibre groups. This pattern was reflected in the effect size analyses, which demonstrated a moderate effect of fibre intake within the TEC cohort (Hedges’ *g* = 0.59), and a negligible effect in the PTSD cohort (Hedges’ *g* = - 0.18). Smaller subgroup sizes meant that the confidence intervals were wide, however these findings suggest that functional butyrate-production pathways may be more sensitive to PTSD-related effects than broader taxonomic groups. Furthermore, this raises the possibility that individuals living with PTSD symptoms may not be getting the same potential microbial benefit from a high fibre diet as others not living with PTSD.

The first study parameter investigated was self-reported mental health symptoms. As hypothesised, participants who met PTSD diagnostic criteria demonstrated a significantly higher symptom load across all self-report mental health scales. This indicated that the PTSD study cohort had not only experienced trauma but were living with substantial and recent psychological distress. Participant PCL-5 scores were significantly and positively correlated with DASS Stress subscale scores, indicating that greater PTSD symptom severity was associated with higher levels of recent perceived stress. The strong relationship between PTSD symptom severity and perceived stress is consistent with current understanding of PTSD. PTSD is characterised by persistent dysregulation of stress-related pathways, including the HPA axis (Lawrence & Scofield, 2024; Schumacher et al., 2019). Combined with symptoms such as intrusive thoughts and distressing reminders of traumatic events, this provides a biologically plausible explanation for the elevated stress burden observed in the PTSD cohort.

The second study parameter investigated was cognitive function. Premature and accelerated cognitive decline has been extensively reported in older adults with PTSD (Prieto et al., 2023; Sumner et al., 2017) and has been linked to an increased risk of dementia, including Alzheimer’s disease (Flatt et al., 2018). Participants with PTSD generally performed worse on selected CANTAB outcomes, although not all cognitive domains were equally affected. This suggests that some aspects of cognition may be more susceptible to the effects of PTSD symptom severity than others. The PTSD group performed worse in Spatial Working Memory Strategy (SWMS). Most notably, higher PCL-5 scores were significantly and positively correlated with worse SWMS scores (*p* < 0.05). Multiple linear regression analyses demonstrated that, after adjustment for age, PCL-5 scores remained significantly associated with SWMS performance (*β* = 0.057, *p* = 0.011), whereas age was not independently associated with SWMS performance (*p* = 0.887). These findings support previous literature suggesting that PTSD symptom severity may predict cognitive decline more accurately than traditional AD biomarkers such as beta-amyloid (Aβ) and tau (Prieto et al., 2023; Van Rossum et al., 2025). Furthermore, Paired Associates Learning (Total Errors Adjusted), which assesses new and visual learning, was significantly worse in the PTSD group (*p* < 0.05). Research has indicated that those with PTSD have worse recall than those without PTSD, which is consistent with the results of this study (Golier et al., 2003). Rapid Visual Processing was also significantly different between groups, with PTSD participants having slower reaction times (*p* < 0.05). When Spearman’s correlation analyses were conducted between PCL-5 scores and age, higher RVPMDL scores were associated both with age (r_s_ = 0.55, *p* < 0.05) and PCL-5 scores (r_s_ = 0.4, *p* < 0.05). Multiple linear regression analyses further demonstrated that both age (*β* = 6.210, *p* = 0.007) and PCL-5 (*β* = 1.883, *p* = 0.008) scores were independently and significantly associated with RVPMDL performance.

While correlations between PCL-5 scores, cognition, and mental health status were clear, microbiome data presented challenges when determining trends and statistical significance, likely due to the substantial variability between individuals in this study cohort, combined with the relatively small sample size. The high level of inter-individual variability observed in the gut microbiome meant that biologically meaningful trends may not always have reached statistical significance, particularly in smaller cohorts. There were no significant phylum-level differences between the study groups, highlighting the importance of considering functional pathways and quantifying butyrogenic bacteria when investigating gut microbiome function. Conclusions regarding differences in butyrate-producing pathways between PTSD and TEC participants were limited, as most targets did not differ significantly between groups. However, *buk* demonstrated the strongest evidence of group-level differences. When participants were stratified by fibre intake, differences in *buk* abundance approached statistical significance (H(3) = 7.613, *p* = 0.055), and *buk* relative abundance was significantly lower in PTSD participants than TEC participants (W = 63, *p* = 0.024). These findings suggest that functional butyrate-production pathways may be more sensitive to PTSD-related differences than broader taxonomic composition. Furthermore, higher PTSD symptom severity was associated with lower log_10_-transformed *buk* abundance, and this association remained significant after accounting for dietary fibre intake. Most notably, despite previous literature findings suggesting strong associations between fibre intake and butyrogenic bacterial abundance (Daskova et al., 2021; Liu et al., 2018), fibre intake was not independently associated with the latter or *buk* abundance in this study. This suggests that PTSD symptom burden may be more closely related to variation in *buk* abundance than dietary fibre intake in this cohort.

These findings raise an important consideration for older adults, where the composition of the gut may already be altered by ageing, comorbid gastrointestinal conditions, or other chronic physical and mental health conditions. In these populations, overall fibre intake may not accurately capture factors that influence butyrate production. The abundance of the butyrogenic community may instead be influenced more by the characteristics of the fibre consumed than by total fibre intake alone. Factors such as fibre type (e.g. soluble versus insoluble fibre) and fermentability may play a more important role in determining butyrate production and butyrogenic bacterial abundance (McIntyre et al., 1993). This warrants further investigation to better understand how specific dietary characteristics can modify butyrogenic bacteria in older adults with chronic mental health symptoms.

The disassociation between dietary fibre intake and *buk* abundance warrants further consideration. Previous research has suggested that *buk* and *but* pathways may respond differently under disease conditions (namely, ulcerative colitis) (Vital et al., 2013), indicating these genes may capture different aspects of microbial metabolism. Furthermore, differences in butyrogenic pathway abundance have been observed across animal species with contrasting dietary patterns (i.e. more *buk* in carnivores, almost absent in herbivores) (Vital et al., 2015), suggesting that broader dietary influences may play a greater role in pathway abundance than fibre alone.

A further alternative explanation is that the relationship between dietary fibre and butyrogenic bacteria may be influenced by stress-related physiological processes. Previous findings have implicated differences in microbial composition in conditions such as major depressive disorder (Radjabzadeh et al., 2022), PTSD (Hemmings et al., 2017), and anxiety (Jiang et al., 2018). Trauma-exposed controls exhibited marked differences in butyrate-gene abundance according to fibre intake level, while the PTSD fibre-stratified groups were comparatively similar, even if participants had significantly different fibre intakes (e.g. 10 g/day vs 40 g/day). This was reflected by the moderate effect size of fibre intake in the TEC group, compared with the negligible effect in the PTSD group. This pattern supports previous findings, that chronic psychological stress is a modulator of the gut microbiome. Although speculative, this raises the possibility that the microbiome of individuals living with a high symptom burden may not respond to dietary fibre in the same way as individuals with lower symptom burden. Additional research is required to further determine whether PTSD-related physiological changes influence the capacity of dietary fibre interventions alone to modify butyrogenic communities.

Notably, this study has recognised limitations that need to be acknowledged. The results of this study should be interpreted as exploratory rather than definitive or diagnostic, largely due to the small sample size. Functional assessments of the gut microbiome should be repeated in larger cohorts to provide a comprehensive assessment of the microbiome’s capacity to produce SCFAs, such as butyrate. This was particularly evident in fibre-stratified analyses, where effect size estimates suggested potentially meaningful differences within the TEC cohort, but confidence intervals remained wide due to the subgroup sample sizes and inter-individual variability. Additionally, dietary intake was estimated using the FibreScreen questionnaire, which focuses primarily on fibre consumption, but does not capture other dietary factors that may influence microbiome composition, such as protein, fat, or other overarching dietary patterns. Furthermore, it was acknowledged that mental health symptoms and dietary habits can fluctuate over time, while survey data were collected up to three months prior to biological sample collection. As such, survey responses may not have fully captured participant status at the time of sampling. It should also be noted that the presence of butyrate-producing genes (as tested in this study) does not necessarily indicate active gene expression or butyrate production at the time of sampling, which should be explored in further studies.

## Conclusions

Overall, this pilot study provides preliminary evidence that PTSD, particularly symptom severity, is associated with greater mental health burden and decreased performance in selected aspects of cognitive function and does impact some aspects of the gut microbiome. Ultimately, the results of this study indicate that older adults living with PTSD experience a persistent and significant burden of stress and trauma-related symptoms. Through engagement with participants throughout this study, it was evident that many individuals had lived with these symptoms for decades. The findings of the GUMP pilot study demonstrated significant differences in cognitive domains associated with executive function, working memory, and learning, thereby supporting existing evidence that PTSD in older adults can contribute to premature cognitive decline. As such, these findings emphasise the importance of holistic approaches to PTSD management, that consider psychological, cognitive, and physiological health, particularly in older adults. Future research should continue to investigate how modifiable factors (such as diet and the gut microbiome) could support resilience and healthy brain ageing in trauma-exposed populations, while developing interventions that reduce the chronic neurological consequences associated with PTSD.

---

**Supplementary Table 1.**
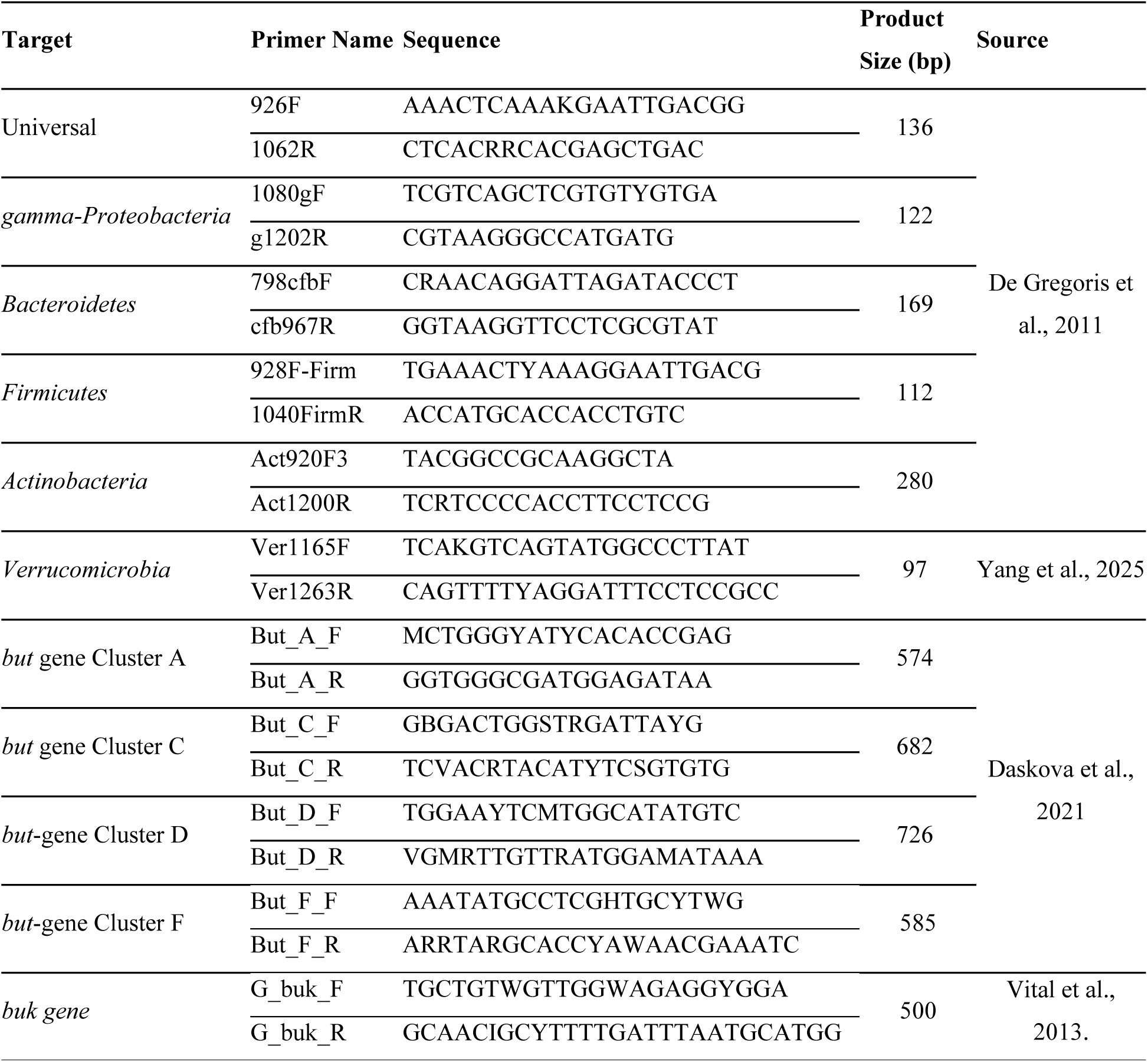
Primers used for qPCR Assays.

**Supplementary Table 2.** Revised Primer Concentrations for GUMP Study.

| Primer | Primer Concentration (µM) | Annealing Temperature (°C) |
| --- | --- | --- |
| Universal (16S gene) | 0.5 | 61.5 |
| *Firmicutes* | 0.4 | 61.5 |
| *Bacteroidetes* | 0.4 | 61.5 |
| *gamma-Proteobacteria* | 0.4 | 61.5 |
| *Actinobacteria* | 0.4 | 61.5 |
| *Verrucomicrobia* | 0.5 | 60 |
| *but-A* | 0.8 | 60 |
| *but -C* | 1.2 | 60 |
| *but -D* | 1.2 | 60 |
| *but -F* | 0.8 | 60 |
| *buk* | 0.83 | 54 |

**Supplementary Table 3.**
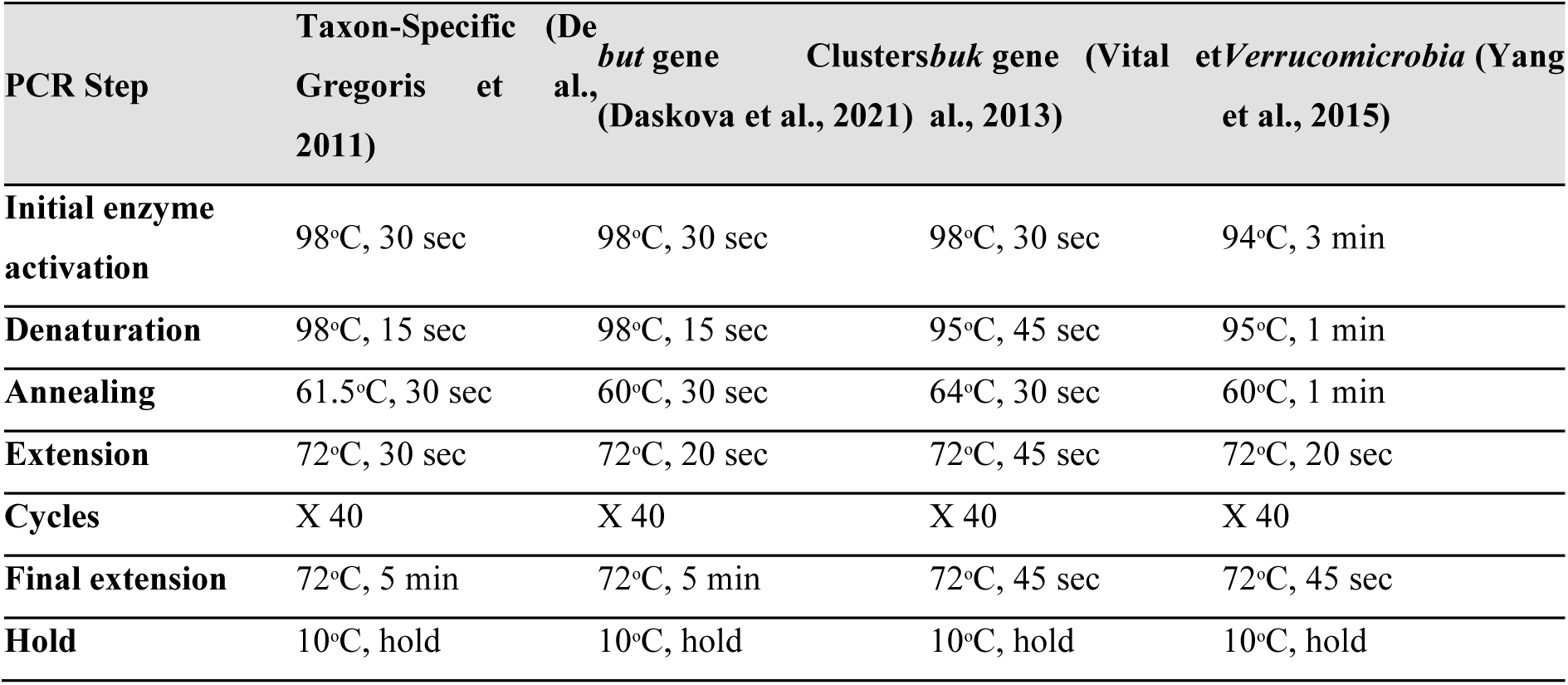
Final Optimised qPCR cycling conditions used for microbiome assays.

**Supplementary Table 4.** Relative abundance (compared to 16S Universal gene total) of major phylum targets.

| Phylum | TEC Low<br>(*n* = 7) | TEC High<br>(*n* = 9) | PTSD Low<br>(*n* = 9) | PTSD High<br>(*n* = 6) | H(3) | *p*-value |
| --- | --- | --- | --- | --- | --- | --- |
| *Actinobacteriota* | 0.9 ± 1.4 | 3.4 ± 5.9 | 5.1 ± 13.4 | 1.7 ± 2.7 | 0.83 | 0.841 |
| *Bacteroidota* | 7.6 ± 5.9 | 14.8 ± 20.6 | 8.4 ± 10.9 | 7.6 ± 8.9 | 1.71 | 0.635 |
| *Firmicutes* | 34.8 ± 18.9 | 56.6 ± 36.8 | 21.4 ± 17.3 | 30.0 ± 18.4 | 1.67 | 0.644 |
| *gamma-Proteobacteria* | 0.1 ± 0.2 | 0.4 ± 0.6 | 0.2 ± 0.3 | 0.1 ± 0.1 | 0.64 | 0.888 |
| *Verrucomicrobiota* | 0.003 ± 0.01 | 1.4 ± 3.8 | 1.9 ± 3.4 | 0.3 ± 0.4 | 4.34 | 0.227 |

Note. All phylum-level relative abundance data violated assumptions of normality according to the Shapiro-Wilk test ( $p < 0.05$ ). Subsequently, Kruskal-Wallis tests were used to compare groups. Values are presented as mean ± standard deviation. H(3) is the Kruskal-Wallis test statistic with three degrees of freedom.

**Supplementary Table 5.** Relative abundance (compared to 16S gene total) of butyrogenic targets.

| Butyrate<br>Gene Target | TEC (*n* = 16) Mean ±<br>SD | PTSD (*n* = 15) Mean ±<br>SD | W | *p*-value |
| --- | --- | --- | --- | --- |
| *but-A* | 0.00005 ± 0.00011 | 0.00016 ± 0.00052 | 123 | 0.922 |
| *but-C* | 0.00310 ± 0.00488 | 0.00187 ± 0.00151 | 117 | 0.922 |
| *but-D* | 0.00455 ± 0.01410 | 0.00147 ± 0.00193 | 111 | 0.741 |
| *but-F* | 0.00117 ± 0.00159 | 0.00081 ± 0.00128 | 109 | 0.682 |
| *buk* | 0.00145 ± 0.00438 | 0.00013 ± 0.00027 | 63 | 0.024* |

Note. All butyrate synthesis pathway gene relative abundance data violated assumptions of normality according to the Shapiro-Wilk test ( $p < 0.05$ ). Subsequently, Wilcoxon-rank sum tests were used to compare groups. Values are presented as mean ± standard deviation. W is the Wilcoxon test statistic.

**Supplementary Figure 1.**
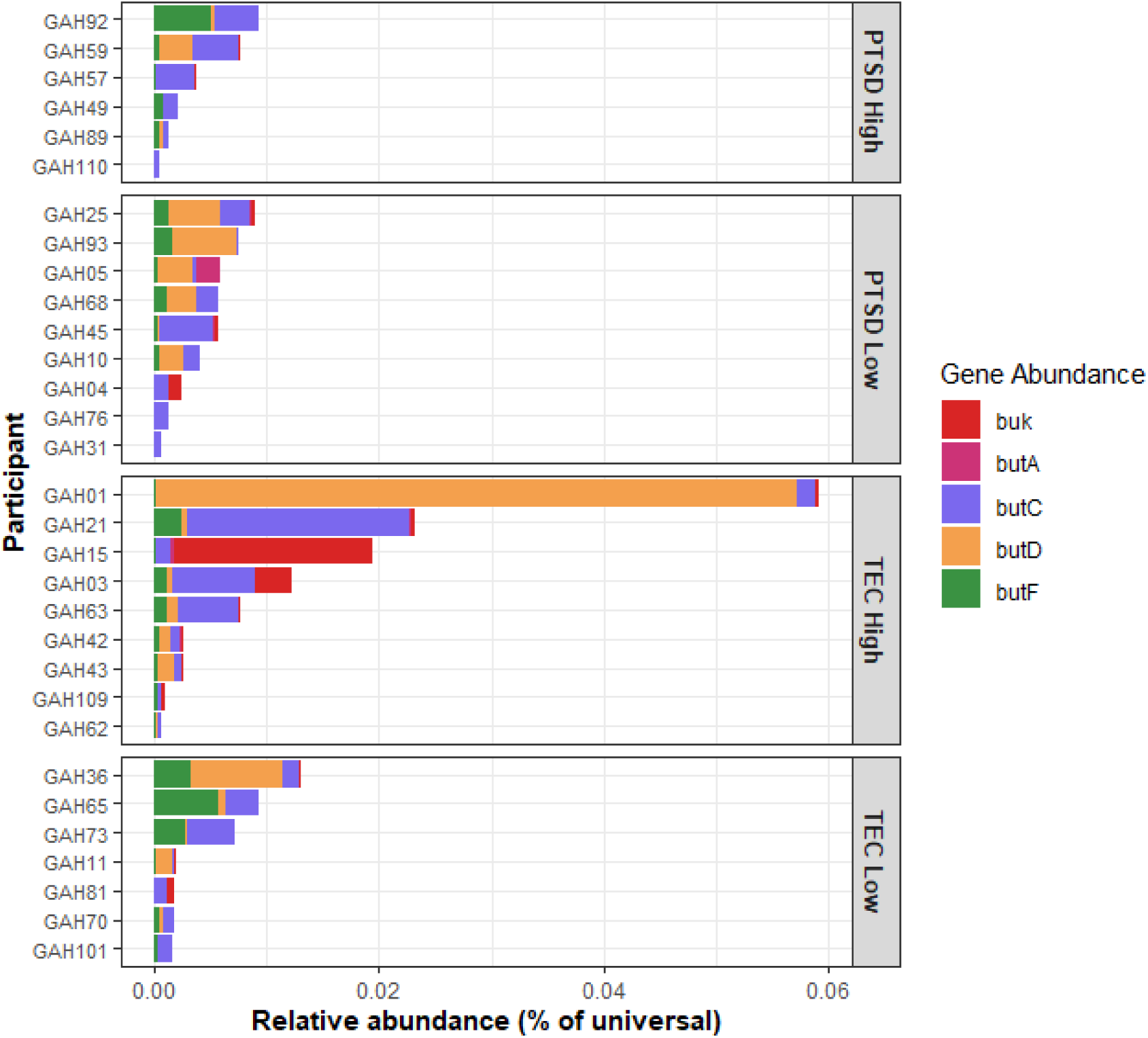
Relative abundance of butyrate-producing genes, calculated as a percentage of the universal 16S rRNA gene, for each individual participant. Substantial between-participant variability was observed, with visual differences in both total butyrate- producing gene abundance and the relative contribution of individual butyrate-producing pathways (*butA, butC, butD, butF*, and *buk*).

## References

American Psychiatric Association. (2013). Diagnostic and Statistical Manual of Mental Disorders (Fifth Edition). American Psychiatric Association. 10.1176/appi.books.9780890425596

Bajaj, J. S. (2019). Alcohol, liver disease and the gut microbiota. Nature Reviews Gastroenterology & Hepatology, 16(4), 235–246. 10.1038/s41575-018-0099-1

Barrett, E. L., Grummitt, L., Jones, S., Rowlinson, K., Vasilopoulos, F., Teesson, M., Mills, K. L., & Sunderland, M. (2026). The prevalence of potentially traumatic events in childhood and associations with mental disorders, suicide and physical health in adulthood: An Australian nationally representative cross-sectional study. Australian & New Zealand Journal of Psychiatry, 60(1), 80–90. 10.1177/00048674251381004

Beristianos, M. H., Yaffe, K., Cohen, B., & Byers, A. L. (2016). PTSD and Risk of Incident Cardiovascular Disease in Aging Veterans. The American Journal of Geriatric Psychiatry, 24(3), 192–200. 10.1016/j.jagp.2014.12.003

Blevins, C. A., Weathers, F. W., Davis, M. T., Witte, T. K., & Domino, J. L. (2015). The Posttraumatic Stress Disorder Checklist for *DSM-5* (PCL-5): Development and Initial Psychometric Evaluation. Journal of Traumatic Stress, 28(6), 489–498. 10.1002/jts.22059

Byers, A. L., Covinsky, K. E., Neylan, T. C., & Yaffe, K. (2014). Chronicity of Posttraumatic Stress Disorder and Risk of Disability in Older Persons. JAMA Psychiatry, 71(5), 540. 10.1001/jamapsychiatry.2014.5

Cunningham, A. L., Stephens, J. W., & Harris, D. A. (2021). Gut microbiota influence in type 2 diabetes mellitus (T2DM). Gut Pathogens, 13(1), 50. 10.1186/s13099-021-00446-0

Daskova, N., Heczkova, M., Modos, I., Videnska, P., Splichalova, P., Pelantova, H., Kuzma, M., Gojda, J., & Cahova, M. (2021). Determination of Butyrate Synthesis Capacity in Gut Microbiota: Quantification of but Gene Abundance by qPCR in Fecal Samples. Biomolecules, 11(9), 1303. 10.3390/biom11091303

Elvers, K. T., Wilson, V. J., Hammond, A., Duncan, L., Huntley, A. L., Hay, A. D., & Van Der Werf, E. T. (2020). Antibiotic-induced changes in the human gut microbiota for the most commonly prescribed antibiotics in primary care in the UK: A systematic review. BMJ Open, 10(9), e035677. 10.1136/bmjopen-2019-035677

Fackelmann, G., Manghi, P., Carlino, N., Heidrich, V., Piccinno, G., Ricci, L., Piperni, E., Arrè, A., Bakker, E., Creedon, A. C., Francis, L., Capdevila Pujol, J., Davies, R., Wolf, J., Bermingham, K. M., Berry, S. E., Spector, T. D., Asnicar, F., & Segata, N. (2025). Gut microbiome signatures of vegan, vegetarian and omnivore diets and associated health outcomes across 21,561 individuals. Nature Microbiology, 10(1), 41–52. 10.1038/s41564-024-01870-z

Fusco, W., Lorenzo, M. B., Cintoni, M., Porcari, S., Rinninella, E., Kaitsas, F., Lener, E., Mele, M. C., Gasbarrini, A., Collado, M. C., Cammarota, G., & Ianiro, G. (2023). Short-Chain Fatty-Acid-Producing Bacteria: Key Components of the Human Gut Microbiota. Nutrients, 15(9), 2211. 10.3390/nu15092211

Geier, T. J., Atkinson, S. N., Pan, A. Y., Mantz-Wichman, M., Jazinski-Chambers, K., Hillard, C. J., & deRoon-Cassini, T. A. (2024). Differences in intestinal bacteria in traumatic injury survivors with and without probable posttraumatic stress disorder. Journal of Affective Disorders, 361, 528–535. 10.1016/j.jad.2024.06.075

Gill, S. K., Rossi, M., Bajka, B., & Whelan, K. (2021). Dietary fibre in gastrointestinal health and disease. Nature Reviews Gastroenterology & Hepatology, 18(2), 101–116. 10.1038/s41575-020-00375-4

Hemmings, S. M. J., Malan-Müller, S., Van Den Heuvel, L. L., Demmitt, B. A., Stanislawski, M. A., Smith, D. G., Bohr, A. D., Stamper, C. E., Hyde, E. R., Morton, J. T., Marotz, C. A., Siebler, P. H., Braspenning, M., Van Criekinge, W., Hoisington, A. J., Brenner, L. A., Postolache, T. T., McQueen, M. B., Krauter, K. S., … Lowry, C. A. (2017). The Microbiome in Posttraumatic Stress Disorder and Trauma-Exposed Controls: An Exploratory Study. Psychosomatic Medicine, 79(8), 936–946. 10.1097/PSY.0000000000000512

Jiang, H., Zhang, X., Yu, Z., Zhang, Z., Deng, M., Zhao, J., & Ruan, B. (2018). Altered gut microbiota profile in patients with generalized anxiety disorder. Journal of Psychiatric Research, 104, 130–136. 10.1016/j.jpsychires.2018.07.007

Kessler, R. C., Aguilar-Gaxiola, S., Alonso, J., Benjet, C., Bromet, E. J., Cardoso, G., Degenhardt, L., De Girolamo, G., Dinolova, R. V., Ferry, F., Florescu, S., Gureje, O., Haro, J. M., Huang, Y., Karam, E. G., Kawakami, N., Lee, S., Lepine, J.-P., Levinson, D., … Koenen, K. C. (2017). Trauma and PTSD in the WHO World Mental Health Surveys. European Journal of Psychotraumatology, 8(sup5), 1353383. 10.1080/20008198.2017.1353383

Kessler, R. C., Berglund, P., Demler, O., Jin, R., Merikangas, K. R., & Walters, E. E. (2005). Lifetime Prevalence and Age-of-Onset Distributions of DSM-IV Disorders in the National Comorbidity Survey Replication. Archives of General Psychiatry, 62(6), 593. 10.1001/archpsyc.62.6.593

Leclercq, S., Forsythe, P., & Bienenstock, J. (2016). Posttraumatic Stress Disorder: Does the Gut Microbiome Hold the Key? The Canadian Journal of Psychiatry, 61(4), 204–213. 10.1177/0706743716635535

Liu, H., Wang, J., He, T., Becker, S., Zhang, G., Li, D., & Ma, X. (2018). Butyrate: A Double-Edged Sword for Health? Advances in Nutrition, 9(1), 21–29. 10.1093/advances/nmx009

Lovibond, S. H., & Lovibond, P. F. (2011). Depression Anxiety Stress Scales [Dataset]. 10.1037/t01004-000

Maier, T. V., Lucio, M., Lee, L. H., VerBerkmoes, N. C., Brislawn, C. J., Bernhardt, J., Lamendella, R., McDermott, J. E., Bergeron, N., Heinzmann, S. S., Morton, J. T., González, A., Ackermann, G., Knight, R., Riedel, K., Krauss, R. M., Schmitt-Kopplin, P., & Jansson, J. K. (2017). Impact of Dietary Resistant Starch on the Human Gut Microbiome, Metaproteome, and Metabolome. mBio, 8(5), e01343–17. 10.1128/mBio.01343-17

Malan-Muller, S., Valles-Colomer, M., Foxx, C. L., Vieira-Silva, S., Van Den Heuvel, L. L., Raes, J., Seedat, S., Lowry, C. A., & Hemmings, S. M. J. (2022). Exploring the relationship between the gut microbiome and mental health outcomes in a posttraumatic stress disorder cohort relative to trauma-exposed controls. European Neuropsychopharmacology, 56, 24–38. 10.1016/j.euroneuro.2021.11.009

Marano, G., Lo Giudice, L., Specogna, E., Chisari, L., Brisi, C., Traversi, G., Mazza, O., & Mazza, M. (2026). From Dysbiosis to Distress: The Gut–Brain Connection in Trauma-Related Disorders. Nutrients, 18(3), 530. 10.3390/nu18030530

McIntyre, A., Gibson, P. R., & Young, G. P. (1993). Butyrate production from dietary fibre and protection against large bowel cancer in a rat model. Gut, 34(3), 386–391. 10.1136/gut.34.3.386

Nilaweera, D., Freak-Poli, R., Ritchie, K., Chaudieu, I., Ancelin, M.-L., & Ryan, J. (2020). The long-term consequences of trauma and posttraumatic stress disorder symptoms on later life cognitive function and dementia risk. Psychiatry Research, 294, 113506. 10.1016/j.psychres.2020.113506

Posit Team. (2025). *RStudio: Integrated Development Environment for R* [Computer software]. Posit software, PBC. http://www.posit.co/

Radjabzadeh, D., Bosch, J. A., Uitterlinden, A. G., Zwinderman, A. H., Ikram, M. A., Van Meurs, J. B. J., Luik, A. I., Nieuwdorp, M., Lok, A., Van Duijn, C. M., Kraaij, R., & Amin, N. (2022). Gut microbiome-wide association study of depressive symptoms. Nature Communications, 13(1), 7128. 10.1038/s41467-022-34502-3

Reynolds, K., Pietrzak, R. H., Mackenzie, C. S., Chou, K. L., & Sareen, J. (2016). Post-Traumatic Stress Disorder Across the Adult Lifespan: Findings From a Nationally Representative Survey. The American Journal of Geriatric Psychiatry, 24(1), 81–93. 10.1016/j.jagp.2015.11.001

Rijnaarts, I., De Roos, N., Zoetendal, E. G., De Wit, N., & Witteman, B. J. M. (2021). Development and validation of the FiberScreen: A short questionnaire to screen fibre intake in adults. Journal of Human Nutrition and Dietetics, 34(6), 969–980. 10.1111/jhn.12941

Sankarganesh, P., Bhunia, A., Ganesh Kumar, A., Babu, A. S., Gopukumar, S. T., & Lokesh, E. (2025). Short-chain fatty acids (SCFAs) in gut health: Implications for drug metabolism and therapeutics. Medicine in Microecology, 25, 100139. 10.1016/j.medmic.2025.100139

Singh, V., Lee, G., Son, H., Koh, H., Kim, E. S., Unno, T., & Shin, J.-H. (2023). Butyrate producers, “The Sentinel of Gut”: Their intestinal significance with and beyond butyrate, and prospective use as microbial therapeutics. Frontiers in Microbiology, 13, 1103836. 10.3389/fmicb.2022.1103836

Steinert, C., Hofmann, M., Leichsenring, F., & Kruse, J. (2015). The course of PTSD in naturalistic long-term studies: High variability of outcomes. A systematic review. Nordic Journal of Psychiatry, 69(7), 483–496. 10.3109/08039488.2015.1005023

Taft, T. H., Bedell, A., Craven, M. R., Guadagnoli, L., Quinton, S., & Hanauer, S. B. (2019). Initial Assessment of Post-traumatic Stress in a US Cohort of Inflammatory Bowel Disease Patients. Inflammatory Bowel Diseases, 25(9), 1577–1585. 10.1093/ibd/izz032

Tomova, A., Bukovsky, I., Rembert, E., Yonas, W., Alwarith, J., Barnard, N. D., & Kahleova, H. (2019). The Effects of Vegetarian and Vegan Diets on Gut Microbiota. Frontiers in Nutrition, 6, 47. 10.3389/fnut.2019.00047

Van-Wehle, T., & Vital, M. (2024). Investigating the response of the butyrate production potential to major fibers in dietary intervention studies. Npj Biofilms and Microbiomes, 10(1), 63. 10.1038/s41522-024-00533-5

Vital, M., Gao, J., Rizzo, M., Harrison, T., & Tiedje, J. M. (2015). Diet is a major factor governing the fecal butyrate-producing community structure across *Mammalia* , *Aves* and *Reptilia*. The ISME Journal, 9(4), 832–843. 10.1038/ismej.2014.179

Vital, M., Howe, A. C., & Tiedje, J. M. (2014). Revealing the Bacterial Butyrate Synthesis Pathways by Analyzing (Meta)genomic Data. mBio, 5(2), e00889–14. 10.1128/mBio.00889-14

Vital, M., Penton, C. R., Wang, Q., Young, V. B., Antonopoulos, D. A., Sogin, M. L., Morrison, H. G., Raffals, L., Chang, E. B., Huffnagle, G. B., Schmidt, T. M., Cole, J. R., & Tiedje, J. M. (2013). A gene-targeted approach to investigate the intestinal butyrate-producing bacterialcommunity. Microbiome, 1(1), 8. 10.1186/2049-2618-1-8

Weathers, F. W., Blake, D. D., Schnurr, P. P., Kaloupek, D. G., Marx, B. P., & Keane, T. M. (2013). The Life Events CHecklist for DSM-5 (LEC-5). https://www.ptsd.va.gov/professional/assessment/te-measures/life_events_checklist.asp

Wickham, H. (with Sievert, C.). (2016). ggplot2: Elegant graphics for data analysis (Second edition). Springer international publishing.

Wickham, H., Averick, M., Bryan, J., Chang, W., McGowan, L., François, R., Grolemund, G., Hayes, A., Henry, L., Hester, J., Kuhn, M., Pedersen, T., Miller, E., Bache, S., Müller, K., Ooms, J., Robinson, D., Seidel, D., Spinu, V., … Yutani, H. (2019). Welcome to the Tidyverse. Journal of Open Source Software, 4(43), 1686. 10.21105/joss.01686

Yaffe, K., Vittinghoff, E., Lindquist, K., Barnes, D., Covinsky, K. E., Neylan, T., Kluse, M., & Marmar, C. (2010). Posttraumatic Stress Disorder and Risk of Dementia Among US Veterans. Archives of General Psychiatry, 67(6), 608. 10.1001/archgenpsychiatry.2010.61

Zeamer, A. L., Salive, M.-C., An, X., Beaudoin, F. L., House, S. L., Stevens, J. S., Zeng, D., Neylan, T. C., Clifford, G. D., Linnstaedt, S. D., Rauch, S. L., Storrow, A. B., Lewandowski, C., Musey, P. I., Hendry, P. L., Sheikh, S., Jones, C. W., Punches, B. E., Swor, R. A., … Haran, J. P. (2023). Association between microbiome and the development of adverse posttraumatic neuropsychiatric sequelae after traumatic stress exposure. Translational Psychiatry, 13(1), 354. 10.1038/s41398-023-02643-8

Zigmond, A. S., & Snaith, R. P. (1983). The Hospital Anxiety and Depression Scale. Acta Psychiatrica Scandinavica, 67(6), 361–370. 10.1111/j.1600-0447.1983.tb09716.x

Zohar, J., Amital, D., Cropp, H. D., Cohen-Rappaport, G., Zinger, Y., & Sasson, Y. (2000). Update on the epidemiology, diagnosis, and treatment of posttraumatic stress disorder. Dialogues in Clinical Neuroscience, 2(1), 37–43. 10.31887/DCNS.2000.2.1/jzohar

